# Proteomics Characteristics Reveal the Risk of T1 Colorectal Cancer Metastasis to Lymph Nodes

**DOI:** 10.1101/2022.09.22.508997

**Authors:** Aojia Zhuang, Aobo Zhuang, Zhaoyu Qin, Dexiang Zhu, Li Ren, Ye Wei, Pengyang Zhou, Xuetong Yue, Fuchu He, Jianming Xu, Chen Ding

## Abstract

**Background:** The presence of lymph node metastasis (LNM) affects treatment strategy decisions in T1NxM0 colorectal cancer (CRC), but the currently used clinicopathological-based risk stratification cannot predict LNM accurately. In this study, we established a classifier for predicting LNM in T1 CRC.

**Methods:** We detected proteins in formalin-fixed paraffin-embedded (FFPE) tumor samples from 143 LNM-negative and 78 LNM-positive patients with T1 CRC and revealed changes in molecular and biological pathways by label-free LC-MS/MS. An effective prediction model was built and validated in a training cohort (N=132) and two validation cohorts (VC1, N=42; VC2, N=47) by machine learning. We further built a simplified classifier with 9 proteins. The expression patterns of 13 proteins were confirmed by immunohistochemistry, and the IHC score of 5 proteins were used to build a IHC predict model.

**Result:** Patients with or without LNM have different molecular signatures. The 55-proteins prediction model achieved an impressive AUC of 1.00 in the training cohort, 0.96 in VC1 and 0.93 in VC2. The 9-protein classifier achieved an AUC of 0.824, and the calibration plot was excellent. We found that 5 biomarkers could predict LNM by the IHC score, with an AUC of 0.825. RHOT2 silence significantly enhanced migration and invasion of colon cancer cells.

**Conclusions:** Our study explored the mechanism of metastasis in T1 CRC and can be used to facilitate the individualized prediction of LNM in patients with T1 CRC, which may provide a guidance for clinical practice in T1 CRC.

## Introduction

Colorectal cancer (CRC) is the third most common cancer worldwide and the third leading cause of cancer-related deaths in Western countries [1–3]. With the introduction of population-based screening programs, a growing number of early invasive CRCs (T1 CRCs) are being diagnosed [4]. It is estimated that the endoscopic removal of adenomatous polyps can reduce CRC-related mortality by more than 50%. Evidence suggests that ESD (endoscopic submucosal dissection) alone is an effective option for T1 CRC patients, who are at low risk for developing lymph node metastasis (LNM), while more extensive radical surgery after ESD is needed for only high-risk patients [5, 6]. According to the current clinical treatment guidelines, which rely on a histopathological examination, approximately 70 to 80% of patients are classified as high risk (poor tumor differentiation, lymphatic/vascular invasion and depth of submucosal invasion >1000 mm). However, the LN status of only 8% to 16% of patients can be accurately predicted by these guidelines; thus, a large number of LNM-negative patients routinely undergo unnecessary additional surgeries, with an associated postoperative mortality rate of 3%-6% [7, 8]. The contradiction between the high rate of additional surgical resection and the fact that only a few people have LNM is due to the lack of accurate diagnostic methods. Therefore, there is an urgent need to develop a new method that can effectively determine LNM in T1 CRC.

Previously, Ozawa et al. and Kandimalla et al. used the microRNA (miRNA) and messenger RNA (mRNA) expression dataset of T1 and T2 CRC patients from The Cancer Genome Atlas (TCGA) as the training cohort to build predictive models, achieved area under the ROC curve (AUC) values of 0.74 and 0.84, respectively [9, 10]. After that, wada et al. validated these miRNA and mRNA signatures in the blood samples [11]. These studies unraveled a new paradigm for a more adequate risk assessment and identification of patients who are true candidates for endoscopic treatment or radical surgery. However, these retrospective studies have several limitations, less than 10% of T1 colorectal cancer samples had lymph node metastasis and almost no specimens removed by ESD were included. Most recently, Kudo et al. developed an algorithm to predict LNM in 4073 patients with T1 CRC by clinicopathological characteristics and achieved an AUC of 0.83 [12]. The accurate prediction of LNM in T1 CRC is a crucial but difficult challenge for scientists worldwide.

Proteins, as executors of biological functions, are receiving much research attention. Several recent proteomic studies of CRC defined new protein signatures, molecular subtypes, and metastasis markers [13–16] and revealed differences in tumorigenesis between right- and left-sided CRC [17]. However, these studies focused on advanced-stage CRC rather than early-stage CRC, and proteomic studies of T1NxM0 CRC are still lacking. There was a recent proteomic study using 21 T1 and T2 CRC patients [18]. Here, we assembled three cohorts of patients with or without LNM (a training cohort, an ESD validation cohort and a prospective validation cohort). A quantitative proteomics approach was used to analyze a total of 221 patients. We developed a high-performance prediction model that can help to reduce the numbers of unnecessary additional surgeries and will benefit the majority of patients.

## Methods

### Patient

In this study, two independent cohorts of consecutive patients who visited the General Surgery Department, Zhongshan Hospital, Fudan University (Shanghai, China) at different time periods were enrolled.

The patients consisted of a training cohort and two different validation cohorts. In the training cohort, we initially collected data on 604 patients who underwent surgical resection with LN dissection between June 2008 and October 2019. We then excluded 233 patients in whom fewer than 12 LNs were examined from the LNM-negative group. After matching the two groups for sex and age, we enrolled 132 patients. For the ESD validation cohort (VC1), we studied 42 consecutive ESD samples before additional surgical resection from June 2008 to October 2019, and the LNM status was then examined by LN dissection. The prospective validation cohort (VC2) comprised 47 prospectively enrolled individuals from November 2019 to April 2020. The model was developed and validated in the retrospective cohort and then prospectively tested in the prospective cohort. In all cohorts, the inclusion criteria (as follows) were the same: aged between 18 and 80 years; and pathological confirmation of colorectal adenocarcinoma, mucinous adenocarcinoma, or signet-ring cell carcinoma with pT1 disease according to the AJCC/UICC TNM staging system, 8th edition. The exclusion criteria (as follows) were also the same in all cohorts: patients who had undergone only endoscopic treatment; those diagnosed with familial adenomatous polyposis, Lynch syndrome, or ulcerative colitis; those who had undergone transanal endoscopic microsurgery; those who developed synchronous invasive carcinomas; and those with missing data. Patients who had received preoperative chemotherapy or radiotherapy were excluded.

### Sample preparation

The FFPE samples derived from 221 T1 CRC patients were collected, and the tumor regions were determined by pathological examination. For clinical sample preparation, sections (10 μm thick) from FFPE blocks were macro-dissected, deparaffinized with xylene, and washed with ethanol. The ethanol was removed completely and the sections were left to air-dry. For this purpose, a hematoxylin-stained section of the same tumor was used as reference. Areas containing 80% or more tumor were examined by pathologist.

Lysis buffer [0.1 M Tris-HCl (pH 8.0), 0.1 M DTT (Sigma, 43815), 1 mM PMSF (Amresco, M145)] was added to the extracted tissues, and subsequently sonicated for 1 min (3 s on and 3 s off, amplitude 25%) on ice. The supernatants were collected, and the protein concentration was determined using the Bradford assay. The extracted tissues were then lysed with 4% sodium dodecyl sulfate (SDS) and kept for 2.5 h at 99 °C with shaking at 1800 rpm. The solution was collected by centrifugation at 12,000 ×g for 5 min. A 4-fold volume of acetone was added to the supernatant and kept in -20 °C overnight. Subsequently, the acetone-precipitated proteins were washed three times with cooled acetone. Filter-aided sample preparation (FASP) procedure was used for protein digestion [19]. The proteins were resuspended in 200 μL 8 M urea (pH 8.0) and loaded in 30 kD Microcon filter tubes (Sartorius) and centrifuged at 12,000 g for 20 min. The precipitate in the filter was washed three times by adding 200 μL 50 mM NH4HCO3. The precipitate was resuspended in 50 μL 50 mM NH4HCO3. Protein samples underwent trypsin digestion (enzyme-to-substrate ratio of 1:50 at 37oC for 18– 20 h) in the filter, and then were collected by centrifugation at 12,000 g for 15 min. Additional washing, twice with 200 μL of MS water, was essential to obtain greater yields. Finally, the centrifugate was pumped out using the AQ model Vacuum concentrator (Eppendorf, Germany).

### Mass spectrometry analysis

Peptide samples were analyzed on a Q Exactive HF-X Hybrid Quadrupole-Orbitrap Mass Spectrometer (Thermo Fisher Scientific, Rockford, IL, USA) coupled with a high-performance liquid chromatography system (EASY nLC 1200, Thermo Fisher). Peptides, re-dissolved in Solvent A (0.1% formic acid in water), were loaded onto a 2-cm self-packed trap column (100-μm inner diameter, 3-μm ReproSil-Pur C18-AQ beads, Dr. Maisch GmbH) using Solvent A, and separated on a 150-μm-inner-diameter column with a length of 15 cm (1.9-μm ReproSil-Pur C18-AQ beads, Dr. Maisch GmbH) over a 75 min gradient (Solvent A: 0.1 % formic acid in water; Solvent B: 0.1 % formic acid in 80 % ACN) at a constant flow rate of 600 nL/min (0–75 min, 0 min, 4% B; 0– 10 min, 4–15% B; 10–60 min, 15–30% B; 60–69 min, 30–50% B; 69–70 min, 50–100% B; 70–75 min, 100% B). The eluted peptides were ionized under 2 kV and introduced into the mass spectrometer. MS was operated under a data-dependent acquisition mode. For the MS1 Spectra full scan, ions with m/z ranging from 300 to 1,400 were acquired by Orbitrap mass analyzer at a high resolution of 120,000. The automatic gain control (AGC) target value was set as 3E+06. The maximal ion injection time was 80 ms. MS2 Spectra acquisition was performed in top-speed mode. Precursor ions were selected and fragmented with higher energy collision dissociation with a normalized collision energy of 27%. Fragment ions were analyzed using an ion trap mass analyzer with an AGC target value of 5E+04, with a maximal ion injection time of 20 ms. Peptides that triggered MS/MS scans were dynamically excluded from further MS/MS scans for 12 s. A single-run measurement was kept for 75 min. All data were acquired using Xcalibur software (Thermo Scientific)

### Peptide and protein identification

MS raw files were processed using the Firmiana proteomics workstation [20]. Briefly, raw files were searched against the NCBI human Refseq protein database (released on 04-07-2013; 32,015 entries) using the Mascot search engine (version 2.3, Matrix Science Inc). The mass tolerances were: 20 ppm for precursor and 50 mmu for product ions collected by Q Exactive HF-X. Up to two missed cleavages were allowed. The database searching considered cysteine carbamidomethylation as a fixed modification, and N-acetylation, and oxidation of methionine as variable modifications. Precursor ion score charges were limited to +2, +3, and +4. For the quality control of protein identification, the target-decoy-based strategy was applied to confirm the FDR of both peptide and protein, which was lower than 1 %. Percolator was used to obtain the quality value (q-value), validating the FDR (measured by the decoy hits) of every peptide-spectrum match (PSM), which was lower than 1 %. Subsequently, all the peptides shorter than seven amino acids were removed. The cutoff ion score for peptide identification was 20. All the PSMs in all fractions were combined to comply with a stringent protein quality control strategy. We employed the parsimony principle and dynamically increased the q-values of both target and decoy peptide sequences until the corresponding protein FDR was less than 1 %. Finally, to reduce the false positive rate, the proteins with at least one unique peptide were selected for further investigation.

### Label-free-based MS quantification of proteins

The one-stop proteomic cloud platform “Firmiana” was further employed for protein quantification. Identification results and the raw data from the mzXML file were loaded. Then for each identified peptide, the extracted-ion chromatogram (XIC) was extracted by searching against the MS1 based on its identification information, and the abundance was estimated by calculating the area under the extracted XIC curve. For protein abundance calculation, the nonredundant peptide list was used to assemble proteins following the parsimony principle. The protein abundance was estimated using a traditional label-free, intensity-based absolute quantification (iBAQ) algorithm [21], which divided the protein abundance (derived from identified peptides’ intensities) by the number of theoretically observable peptides. We built a dynamic regression function based on the commonly identified peptides in tumor samples. According to correlation value R2, Firmiana chose linear or quadratic functions for regression to calculate the retention time (RT) of corresponding hidden peptides, and to check the existence of the XIC based on the m/z and calculated RT. Subsequently, the fraction of total (FOT), a relative quantification value was defined as a protein’s iBAQ divided by the total iBAQ of all identified proteins in one experiment, and was calculated as the normalized abundance of a particular protein among experiments. Finally, the FOT was further multiplied by 105 for ease of presentation, and FOTs less than 10-5 were replaced with 10-5 to adjust extremely small values [22].

### Immunohistochemistry (IHC) staining

FFPE tumor blocks were obtained from the Institute of Pathology at the Affiliated Zhongshan Hospital of Fudan University. Tumor blocks were cut into 4 μm sections. Nonspecific background staining was blocked via a serum-free protein blocker (BOSTER, USA) for 10 min at room temperature. Next, Sections were incubated with anti-SHMT1 (SignalwayAntibody, 31314), anti-PAAF1 (Proteintech, 17650-1-AP), anti-VRK2 (SignalwayAntibody, 43825), anti-ABI1 (SignalwayAntibody, 36723), anti-RHOT2 (Proteintech, 11237-1-AP), anti-SWAP70 (SignalwayAntibody, 42812), anti-TTC19 (Proteintech, 20875-1-AP), anti-ZG16 (Proteintech, 67389-1-Ig), anti-ATAD2 (Cell Signaling Technology, 78568S), anti-BAIAP2 (Proteintech, 11087-2-AP), anti-ISLR2 (Novus Biologicals, AF4526-SP), anti-ITPR2 (SignalwayAntibody, 37666) overnight at 4°C after blocking for one hour at room temperature, according to the manufacturers’ instructions. Then, TMA sections were incubated with secondary biotinylated goat anti-Rabbit/Mouse antibody. For signal detection, samples were incubated with DAB (BOSTER, USA) for 10 min. All specimens were counterstained with hematoxylin and Scott’s blue. Washing steps were conducted with tris-buffered saline with 0.1% Tween (pH 7.4). The IHC staining results were evaluated independently by two pathologists who were blinded to the clinicopathologic data. According to the proportion of positive cells, samples were scored as follows: 0+, none; 1+, <25%; 2+, 25%–50%; 3+, 51%–75%; and 4+, 75%–100%. The staining intensity was evaluated as follows: 0, none; 1, weak; 2, medium; and 3, strong. The final score (range 0–12) was calculated by multiplying the two sub-scores, and divided in to 4 groups: high (IHC score: 9-12), medium (IHC score: 5-8), low (IHC score: 1-4) and ND (IHC score: 0, not detected).

### Statistical analysis

Statistical details of experiments and analyses can be found in the figure legends and main text above. All statistical tests and calculations were performed using SPSS, or R. Protein intensities were log2-transformed for further analysis, apart from coefficient analysis. Statistical significance tests, Chi-square test (SPSS), Wilcoxon rank-sum test (R), as denoted in each analysis. The statistical significance was considered when P value < 0.05. Kaplan–Meier plots (Log-rank test) were used to describe overall survival.

### LNM Prediction Model

The least absolute shrinkage and selection operator (LASSO) method, which is suitable for the regression of high-dimensional data was used to select the most useful predictive proteins from the training cohort. Lasso binary logistic regression was done using the “glmnet” package. A LNM score calculated for each patient via a linear combination of selected features that were weighted by their respective coefficients. To obtain an unbiased estimate of the prediction power of signatures, we performed a 10-fold cross-validation on the training cohort used Logistic Regression. The predictive power was also validated in two validation cohorts. Receiver operating characteristic (ROC) curves were constructed using the pROC R package in the R software v3.5.1, and the area under the ROC curve (AUC) was used to evaluate the diagnostic performance of the selected proteins.

Ninet proteins were further selected to build a simplified classifier using Logistic Regression. Calibration curves were plotted to assess the calibration of the classifier used rms and ResourceSelection R packages in the R software, accompanied with bootstrapping validation (1,000 bootstrap resamples). Decision curve analysis was conducted to determine the clinical usefulness of the simplified model by quantifying the net benefits at different threshold probabilities in the validation dataset. The decision curve was constructed using the rmda R package in the R software.

The prediction power of IHC scores in single IHC samples were built by SPSS, and the 5 proteins predict model were built by R using Logistic Regression.

### Cell Culture, Transfection, and Cell Lines

We used human colon cancer cell lines SW480, HT29, HCT-116, RKO, DLD1 and LoVo (from General Surgery, Zhongshan Hospital) in this study. All cell lines were cultured in DMEM (Corning Costar) with 10 % FBS (Gibco), 100 units of penicillin and 100 mg/mL streptomycin (Gibco) at 37 °C in 5% CO2. Cells were transfected with siRNAs (si-RHOT2#1: sense (5’-3’), GCUCAACGCUUUCCAGAAATT; antisense (5’-3’), UUUCUGGAAAGCGUUGAGCTT; si-RHOT2#2: sense(5’-3’), GCGUCUACAAGCACCAUUATT; antisense (5’-3’), UAAUGGUGCUUGUAGACGCTT) using Lipofectamine 2000 according to the manufacturer’s protocol. Knockdown efficiency was verified by western blotting.

### Cell migration analysis

For the transwell migration assay, 4 × 104 serum-starved cells were trypsinized and plated onto an FN-coated upper chamber membrane (8-μm pore filter, Corning Costar, Cat. No. 3422) of a transwell chamber with the corresponding inhibitors. The lower transwell chamber was filled with 2.5 % serum containing DMEM. After incubation for 48 h, the filters were removed, and the cells on the membrane were fixed with methanol. The migrated cells on the underside of the membrane were stained with 0.5 % crystal violet. The dye was washed with water, and the cells were examined by microscopy.

## Results

### Cohort Characteristics and Research Design

To identify LNM mechanisms and protein signatures for T1NxM0 CRC, we performed mass spectrometry (MS)-based proteomics to analyze formalin-fixed paraffin-embedded (FFPE) tumor samples from 143 LNM-negative and 78 LNM-positive patients with T1 CRC, totaling 221 individuals. The patients consisted of a training cohort and two different validation cohorts (Figure 1A). The clinicopathologic characteristics of the patients are listed in Figure 1B and Figure 1-source data 1). As previously reported in T1NxM0 CRC, LNM is significantly associated with the submucosal invasion depth (p=0.014, chi-square test), lymphatic or vascular invasion (p<0.001, chi-square test) and poor differentiation (p=0.004, chi-square test) (Figure 1-source data 1). In our cohort, the rate of LNM was related to tumor location, and a significant difference was found between left- and right-sided tumors (p=0.036, chi-square test) (Figure 1- figure supplement 1).

**Figure 1.**
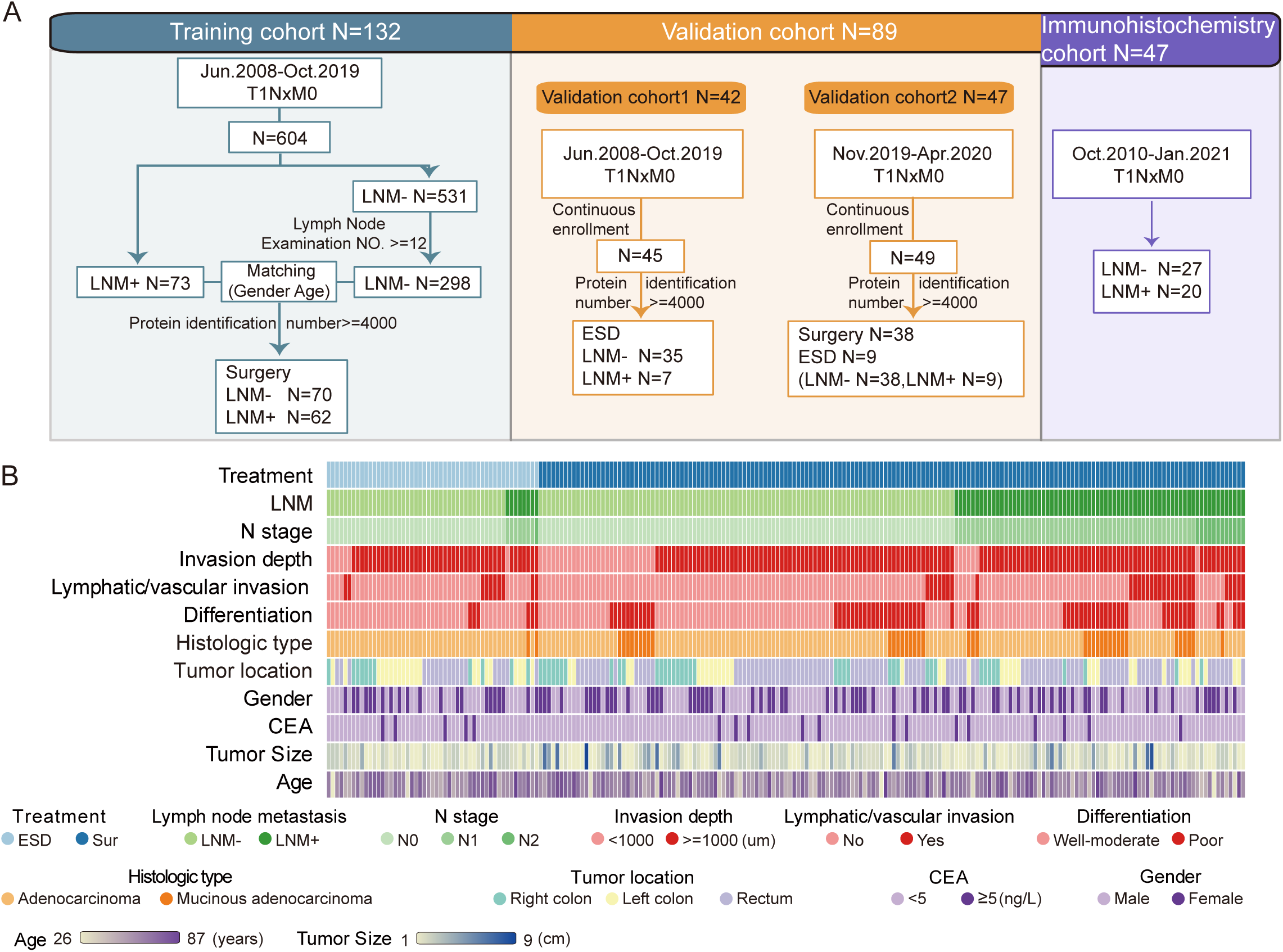
Proteomics landscape of T1 CRC with or without LNM. A. In total, 221 samples were divided into three cohorts for proteomic study: a training cohort (N=132), validation cohort 1 (N=42) and validation cohort 2 (N=47); 47 samples were used for IHC staining. B. The study included 143 LNM-negative 78 LNM-positive patients with T1 CRC and 51 and 170 patients treated with endoscopic submucosal dissection (ESD) or surgical resection, respectively. Clinical parameters are shown in the heatmap. Also see Figure 1- figure supplement 1.

For the proteomic analysis, proteins were obtained from FFPE tumor samples after cleavage with trypsin and analyzed by high-resolution LC-MS/MS on a Q Exactive HF-X mass spectrometer using a label-free technique. Overall, we produced a high-quality dataset (Figure 1- figure supplement 2), more than 13,000 protein groups (with a 1% false discovery rate [FDR] at the peptide and protein levels) were identified, and the identification number of each sample was over 4000 proteins (Figure 1-source data 2). For the bioinformatic analysis, we further filtered the data, as shown in Figure 1- figure supplement 2H (Figure 1-source data 3). Our study represents the first proteomic study of T1 CRC and provides a reliable data source for future research.

### Proteomic Characteristics of the LNM-Negative and LNM-Positive Groups

To identify protein signatures and pathways associated with LNM in T1NxM0 CRC, we investigated the differential proteomic patterns between T1 CRC patients with or without LNM. We comprehensively compared protein signatures and biological differences between the LNM-negative and LNM-positive patients with T1 CRC. We found that 82 and 84 proteins were significantly differentially expressed in LNM-negative and LNM-positive patients, respectively (identified in at least 30% of samples with a log2-fold change [log2FC] > 1 and p<0.05, Wilcoxon rank-sum test) (Figure 2A and Figure 2-source data1). To search for druggable targets in LNM-positive T1 CRC, we investigated 84 proteins that were overrepresented in LNM-positive patients and identified 19 US Food and Drug Administration (FDA)-approved drug targets: CBR3, CES2, CHP1, F13A1, GBA, GNG2, GPRC5A, GUCY1B3, PTGIS, PTK2B, RAB9A, RBP1, RRM2, SLC5A1, SQLE, STK3, STK4, VWF, and WNK1 (Figure 2B and Figure 2-source data2). Furthermore, 34 of these proteins, including ATAD2, BAIAP2, CEACAM6, FARS2, MX2, OSBPL5, SERPINB5, SERPINB8, SHC1, UBE2Z, YAP1, and ZG16, were identified as potential drug targets (Figures 2B) [23]. These potential druggable markers may provide insights into new precision medicine for T1NxM0 CRC and further benefit targeted treatment.

**Figure 2.**
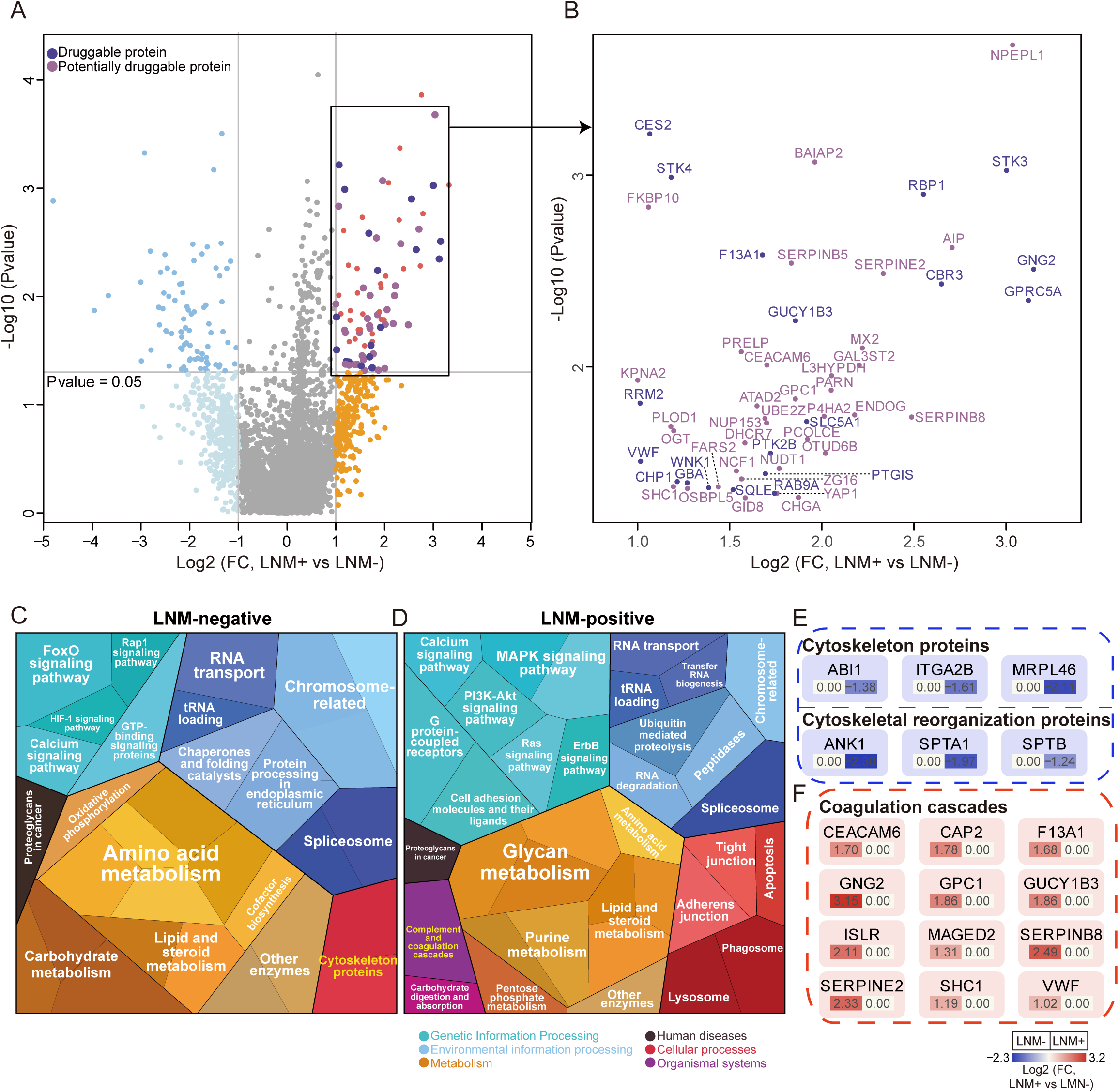
Protein signatures and functional differences between LNM-negative and LNM-positive patients with T1 CRC. A. A volcano plot showing proteins overexpressed in LNM-negative or LNM-positive patients (light blue and orange indicate proteins found in >30% of samples and a fold change more than 2, whereas blue and red indicate proteins with p < 0.05; other proteins are shown in gray). Dark and light purple represent druggable and potentially druggable proteins based on the Drug Gene Interaction Database (http://www.dgidb.org/). B. A scatterplot showing druggable (dark purple, N=19) and potentially druggable (light purple, N=34) proteins based on the Drug Gene Interaction Database (http://www.dgidb.org/) overexpressed in LNM-positive patients. C&D. Pathway enrichment (KEGG) analysis was performed using Proteomaps (www.proteomaps.net). Upregulated pathways in LNM-negative and LNM-positive patients. E&F. Details of proteins involved in the cytoskeletal remodeling (E) and coagulation cascades (F).

To further explore the biological processes associated with LNM, we constructed proteomaps [24] of the differentially expressed proteins according to their Kyoto Encyclopedia Genes and Genomes (KEGG) pathway annotations (Figure 2C, 2D). Diverse RNA metabolism-related proteins were highly expressed in both groups (EIF4E, INTS4, CELF2, SEL1L, etc., in the LNM-negative group; NUP153, TIAL1, FARS2, TRMT11, etc., in the LNM-positive group), while ubiquitin-mediated proteolysis, an oncogenic pathway, was upregulated in the LNM-positive group. In addition to metabolism-related proteins, the LNM-negative group had higher proportions of amino acid metabolism-related (TMLHE, AUH, SHMT1, etc.), carbohydrate metabolism-related (GLA, GUSB, MAN1B1, etc.) and oxidative phosphorylation-related (COX15, etc.) proteins, whereas the LNM-positive group had higher proportions of glycan metabolism-related proteins (OGT, GAL3ST2, PRELP, etc.). Some tumor metastasis-related signaling pathways, including the MAPK, PI3K- Akt, Ras, and ErbB signaling pathways (STK3, STK4, VWF, GNG2, SHC1, etc.), as well as cellular process categories (tight junction, adherens junction, apoptosis, phagosome, and lysosome), were also enriched in LNM-positive patients.

Cytoskeletal proteins, including SPTA1, SPTB and ANK1, and proteins related to cytoskeletal remodeling (ABI1, MRPL46 and ITGA2B), were down-regulated in patients with LNM compared with those without LNM. The rearrangement of cytoskeletal proteins may also be responsible for LNM in patients with T1 CRC (Figure 2E).

Interestingly, the LNM-positive group was significantly enriched in coagulation cascades, and previous studies have indicated that the inhibition of coagulation greatly limits cancer metastasis [25]. Twelve of the 84 proteins that were elevated in LNM-positive patients were related to coagulation cascades (Figure 2E), and many of these 12 proteins are known or suspected to be linked to CRC or other cancer metastasis. For instance, CEACAM6and SERPINE2 are risk genes for colorectal liver metastases [26, 27]. VWF, SHC1 and CAP2 have been reported to be elevated in LNM-positive patients with gastric cancer [28–30]. Moreover, F13A1 and GPC1are biomarkers for melanoma metastasis [31].

### Characterization of Mucinous Colorectal Adenocarcinoma

In agreement with previous reports [5], we found that LNM was significantly associated with the differentiation status (p=0.004, chi-square test) and histologic type in the T1 CRC cohort (p=0.004, chi-square test) (Figure 1- figure supplement 1C and S1D). More than 50% of patients with mucinous colorectal adenocarcinoma had LNM. Thus, we divided our cohort into three subgroups: those with well to moderately differentiated adenocarcinoma (DS1, N=149), poorly differentiated adenocarcinoma (DS2, N=32) and mucinous adenocarcinoma (DS3, N=40) (p=0.007, chi-square test) (Figure 2- figure supplement 1A and Figure 2-source data1). To identify the molecular characteristics of different groups, we determined the significantly changed proteins in each group (identified in at least 30% across groups with a log2-fold change [log2FC] > 1 and p<0.05, Kruskal-Walli’s test) (Figure 2- figure supplement 1B and Figure 1- figure supplement 2H). As a result, 140 proteins were overexpressed in DS1. Further pathway enrichment analysis revealed that the oxidative phosphorylation and TCA cycle pathways were higher in DS1 than in DS2 and DS3 (p<0.05) Figure 2- figure supplement 1C). In DS2, 178 proteins were highly expressed and enriched in the GTPase activity and Wnt signaling pathways. In DS3, 326 proteins were overexpressed, functioning in more aggressive pathways such as ECM organization, cell migration and vesicle-mediated transport. We also found that DS2 shared some features with the other two groups. For example, a SUMOylation-related pathway was identified in both DS1 and DS2, and both DS2 and DS3 were characterized by elevated levels of the NFkB signaling pathway. In conclusion, different differentiation statuses and histologic types in T1 CRC indicate specific pathways. We then studied the relationship between the overrepresented proteins according to histological type and LNM-related proteins. Interestingly, we found a significant negative correlation in the mucinous adenocarcinoma group (Pearson correlation coefficient=-0.53, p<0.05) but not in the other two groups, indicating that mucinous adenocarcinoma has a unique LNM mechanism (Figure 2- figure supplement 1D). Mucinous adenocarcinoma is a distinct subtype of adenocarcinoma and is characterized by abundant mucinous components. In our data, glycoproteins and related enzymes were overexpressed in the mucinous adenocarcinoma group compared with the nonmucinous adenocarcinoma group (Figure 2- figure supplement 1E), indicating that glycoproteins are an important component of mucinous adenocarcinoma.

To explore the mechanism of LNM in mucinous adenocarcinoma in T1 CRC, we focused on the functions and characterizations of the 326 proteins that were highly expressed in the mucinous adenocarcinoma group. We found that 6 integrins (ITGA1, ITGAV, ITGA11, ITGA9, ITGB3 and ITGB5) and 11 reported extracellular vesicle (EV) markers (ADIRF, CSPG4, DPP4, ENTPD1, LRG1, PECAM1, PLVAP, RAB25, TMEM2, TTR and YBX1) were overexpressed in the mucinous adenocarcinoma group (Figure 2- figure supplement 1F, G). Furthermore, proteins involved in the membrane trafficking and/or vesicle transport pathways, such as ARFGAP1, ARRDC1, and TOR1A. and proteins involved in ECM organization, including DDR1, LAMA5, and TTR, were upregulated in mucinous colorectal adenocarcinoma (Figure 2- figure supplement 1H). Next, the composition of the tumor microenvironment in our cohort was studied using xCell [32] (Figure 2- figure supplement 1I and Figure 2-source data 3). The stromal score was significantly higher in mucinous colorectal adenocarcinoma than in non-mucinous adenocarcinoma (p<0.05, Kruskal-Wallis test), and the signatures of endothelial cells, smooth muscle cells and osteoblasts were enriched in mucinous colorectal adenocarcinoma; however, pericytes were decreased. The increase in endothelial cells, especially lymphatic endothelial cells, and the decrease in pericytes indicated that in mucinous colorectal adenocarcinoma, intratumoral lymphatic vessels rather than blood vessels may increase. Blood vessels are composed of endothelial cells and pericytes [33], and lymphatic vessels are composed of a monolayer of endothelial cells [34]. This suggests that in mucinous colorectal adenocarcinoma, the density of intratumoral lymphatic vessels, rather than blood vessels, is increased, and an increase in intratumoral lymphatic vessel density may correlate with LNM [35]. We employed immunohistochemistry (IHC) to validate observation using commercially available D2-40 (PDPN), a lymphatic endothelial marker, antibodies. Immunostaining showed the D2-40 expression in mucinous adenocarcinoma was significantly higher than that in non-mucinous adenocarcinoma, in agreement with the proteomics data (Figure 2- figure supplement 1J and Figure 2-source data4). In summary, we hypothesize that mucinous adenocarcinoma impacts the tumor microenvironment through EVs, resulting in an increase in intratumoral lymphatic vessel density, thereby promoting LNM (Figure 2- figure supplement 1K).

### Discriminative classifier to Identify T1 CRC with LNM

To discover protein markers that can be used to predict LNM in patients with T1 CRC. We established a classifier could effectively distinguish LNM to guide treatment strategy decisions for T1 CRC. Among 70 LNM-negative and 62 LNM-positive patients in the training cohort, we identified 407 candidate proteins (identified in at least 30% of samples and p<0.1, Wilcoxon rank-sum test in the training cohort) (Figure 1- figure supplement 2H). To determine feature importance (significance of the prediction feature), we employed least absolute shrinkage and selection operator (LASSO) logistic regression and used the LNM status (negative or positive) to determine the discrimination power of each signature (Figure 3- figure supplement 1A and B). We constructed a classifier using the intensities of 55 proteins that facilitated accurate discrimination between LNM-negative and LNM-positive patients with T1 CRC in the training cohort (Figure 3A, Figure 1-source data3 and Figure 3-source data1). The classifier achieved an AUC of 1.00 (95% CI, 1.000) through 10-fold cross-validation in the training cohort, which indicated higher predictive power for LNM than the NCCN guidelines (AUC = 0.561) (p<0.001) (Figure 3B). The Youden’s index-derived cutoff was used as the threshold, and the 55-protein classifier yielded 100% sensitivity and specificity, whereas the NCCN guidelines yielded 93.5% sensitivity and 18.6% specificity. In summary, the 55-protein classifier could be used to more accurately predict the individual probability of LNM in the training cohort (Figure 3B).

**Figure 3.**
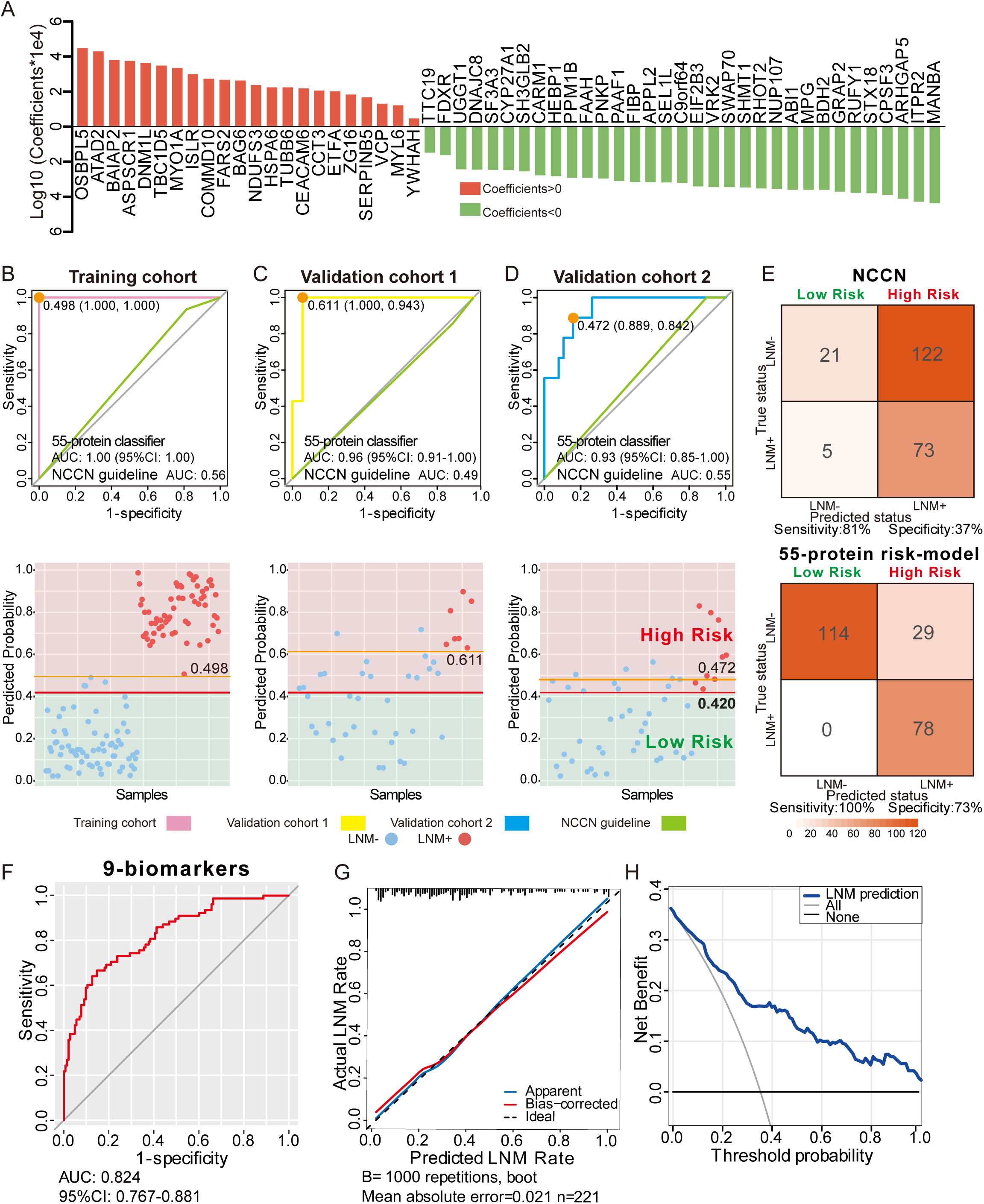
Development and validation of a protein classifier to predict LNM with T1 CRC. A. The predictive relevance of all 55 protein markers to distinguish LNM-positive from LNM-negative T1 CRC patients is represented by a bar chart, and their LASSO coefficients are indicated. Also see Figure 1- figure supplement 2H, Figure 3- figure supplement 1A and B. B, C, D. Top: Receiver operating characteristic (ROC) curve with the area under the curve (AUC) for the protein classifier of the training cohort (B), validation cohort 1 (C) and validation cohort 2 (D). Bottom: Scatterplot representing the score of each patient with (red dot) or without (blue dot) LNM, the optimal threshold (Youden’s index) of each curve (orange line) and the safety cutoff line (red line). E. Classification error matrix using NCCN guidelines and safety cutoff from our 55-protein-model. F, G, H. ROC curve of the optimized 9-biomarker classifier using binary logistic regression (E), calibration curve of the optimized model (F), and cost benefit decision curves (G) in 221 patients.

To validate the predictive power of this classifier, a consecutive dataset from the ESD cohort (VC1, N = 42) was adopted. All the samples in the ESD cohort (35 LNM-negative and 7 LNM-positive individuals) were resected by endoscopy in the clinic. The 55-protein classifier achieved an AUC of 0.96 (95% CI, 0.917 to 1.000) in distinguishing LNM-positive from LNM-negative patients with 100% sensitivity/94.3% specificity in VC1 (Figure 3C). However, the AUC of the NCCN guidelines was 0.49 (85.7% sensitivity/11.4% specificity) (p<0.001) (Figure 3C). This result demonstrates that the proteomic classifier performs better than the NCCN guidelines.

To further assess the value of the proteomic classifier in the clinic, we utilized a prospective validation cohort (VC2) consisting of 47 patients (LNM-negative, N = 38; LNM-positive, N =38); 9 of whom received endoscopic resection, and the others received surgical resection. The classifier based on the intensities of the 55 proteins achieved 88.9% sensitivity and 84.2% specificity in distinguishing LNM-negative and LNM-positive T1 CRC patients, with an AUC of 0.93 (Figure 3D). For the NCCN guidelines, the AUC was 0.55 (p<0.001), which was relatively lower than that of the proteomic indicator.

To ensure the safety of those who have positive LNM, 0.420 was regarded as a cutoff value, when stratifying the patients into “high-risk” and “low-risk” groups by predicting risk score for LNM range from 0 to 1, and at this threshold, all patients in the low-risk group were LNM-negative (Figure 3B-D and Figure 3-source data2). When we used current clinical treatment guidelines (NCCN guidelines), it resulted in stratifying 88% patients (195 of the 221) into a high-risk category and the remaining 12% (26 of 195) into a low-risk group, only 37% (73 of 195) of patients were actually high risk, 63% (122 of 195) of patients were underwent unnecessary additional surgery, and there are 5 (9.2%, 5 of 26) LNM-positive cases mistakenly assigned to the low-risk group (Figure 3E and Figure 1-source data1). In contrast, of the 107 patients who were classified as high risk by our model, 78 were had LNM (72%), indicating that only 27.1% (29 of 104) of the all patients with T1 CRC were overtreated, and all the patient stratified into the low-risk group were LNM-negative (Figure 3E and Figure 3-source data2).

In conclusion, compared to current strategies, our 55-protein classifier can more accurately distinguish LNM-negative and LNM-positive T1 CRC patients to better guide clinical decision-making and determine whether a patient needs additional surgery after ESD.

### Simplified classifier to Identify T1 CRC with LNM

To further reduce the complexity of the indicator, 19 proteins with significant expression changes (log2-fold change [log2FC] > 1 and p<0.05 in 221 samples) were selected from 55 protein markers (SHMT1, PAAF1, VRK2, SEL1L, ITPR2, CPSF3, ABI1, RHOT2, SWAP70, and TTC19 were expressed higher in 143 LNM-negative patients, whereas OSBPL5, FARS2, ZG16, ATAD2, CEACAM6, SERPINB5, COMMD10, BAIAP2, and ISLR were expressed higher in 78 LNM-positive patients), followed by a binary logistic regression analysis that resulted in a final model comprising 9 proteins (ATAD2, CEACAM6, COMMD10, FARS2, ITPR2, RHOT2, SERPINB5, SWAP70, VRK2) (Figure 3-source data3). The 9-protein classifier also demonstrated excellent performance for identifying LNM when we assessed its calibration, discrimination, and clinical usefulness (Figure 3F-H). After using bootstraps with 1,000 resamples for validation, the AUC of the simplified model was 0.824 (95% CI, 0.767 to 0.881) (Figure 3F), and the calibration curve demonstrated good agreement between the predicted status and the true status, with an error of 0.021 (Figure 3G). The decision curve showed that when the threshold probability was > 5%, use of the 9-protein classifier to predict LNM added more benefit than either the treat-all-patients scheme or the treat-none scheme (Figure 3H).

### Five IHC Biomarkers to predict LNM

To further validate the expression patterns of the biomarkers from the proteomic results, 13 proteins from the 19 differently expressed proteins were stained for in LNM-negative and LNM-positive FFPE specimens via IHC. IHC staining was first used in 22 FFPE T1 CRC cases for proteins higher expression in LNM-negative patients, and 21 FFPE T1 CRC cases for proteins higher expression in LNM-positive patients (Figure 4-source data1). For the IHC analysis, protein abundance varied according to the adopted scoring system (on a scale of 0 to 12); thus, we generated a staining scale for all cases (Figure 4A and Figure 4 - figure supplement 1A). In agreement with the proteomics data, staining of ABI1, ITPR2 and RHOT2 showed an overall increase in LNM-negative patients (Figure 4A and B), whereas ATAD2 and ISLR showed an overall increase in LNM-positive patients (Figure 4A and B) according to the proteomics data. However, the protein levels of BAIAP2, CEACAM6, PAAF1, SHMT1, SWAP70, TTC19, VRK2 and ZG16 were not significantly different between LNM-negative and LNM-positive patients (Figure 4- figure supplement 1A).

**Figure 4.**
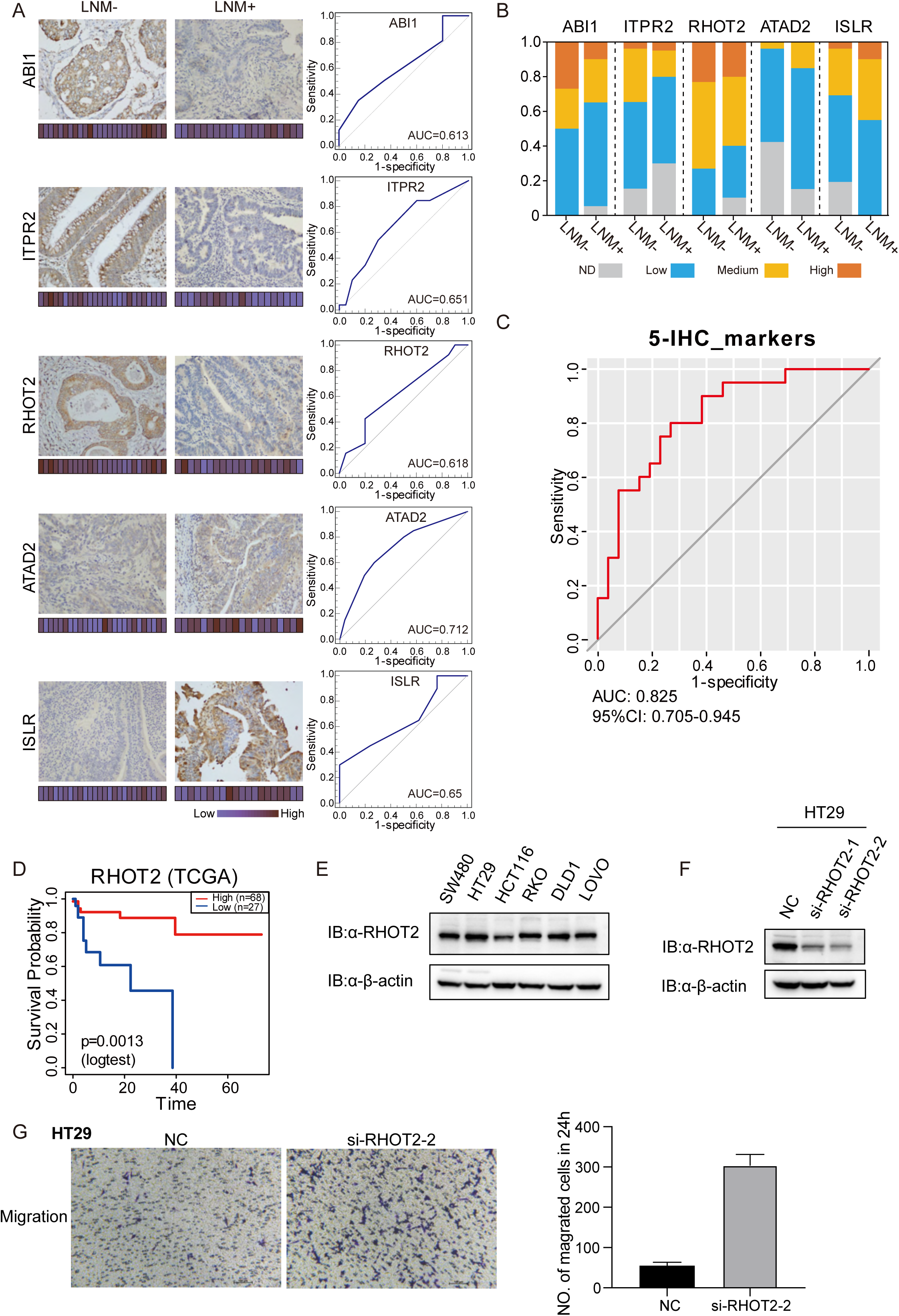
Immunohistochemical staining of targeted proteins. A. T1 CRC samples from a set of 47 cases were used to verify the abundance of ABI1, ITPR2, RHOT2, ATAD2 and ISLR. The scores that represent the sum of the intensities and percentage of protein staining in the LNM-positive or LNM-negative patients are shown as a heat map. (Histological images obtained using a ×40 objective, scale bars, 200 um). The ROC curve of each protein was built by their IHC score. B. Proportions of IHC samples with high (IHC score: 9-12), medium (IHC score: 5-8), or low (IHC score: 1-4) staining. ND: not detected. C. ROC curve of the 5 proteins classifier using IHC score by binary logistic regression. D. The overall survival of patients with colon cancer analyzed on the basis of TCGA database. E. The RHOT2 protein expression in human colon cancer cells (SW480, HT29, HCT-116, RKO, DLD1 and LoVo) measured by western blotting. F. The protein expression of RHOT2 in HT29. G. The migration ability of colon cancer cells was detected by transwell assay.

To further validate the immunohistochemical results, more patients were added. We examined the expression of ABI1, ITPR2, RHOT2, ATAD2 and ISLR in a total of 47 T1 CRC cases (27 LNM-negative patients and 20 in LNM-positive patients). We then evaluated the predictive power of individual proteins to distinguish patients with and without LNM by the AUC. All 5 differentially expressed proteins, namely, ABI1 (AUC=0.613), ITPR2 (AUC=0.651), RHOT2 (AUC=0.618), ATAD2 (AUC=0.712) and ISLR (AUC=0.65), showed good discrimination according to their IHC scores (Figure 4A). Using binary logistic regression to analyze the results of these proteins, we obtained a 5-protein classifier using IHC score. The 5-protein IHC classifier achieved an AUC of 0.825 in 47 patients.

### RHOT2 promotes migration and invasion of colon cancer cells

It has previously been reported that low levels of ABI1 and high levels of ADAT2 or ISLR, result in an increase in extracellular matrix (ECM) degradation, migration, and cell invasion in colon cancer. However, whether RHOT2 could affect the LNM in colon cancer remains unclear. As shown in Figure 4D, the analysis on the basis of the TCGA database suggested that the low level of RHOT2 is related to the low overall survival of patients with colon cancer (Figure 4D, P < 0.05) [14].

Then, we detected the RHOT2 expression in human colon cancer cells (SW480, HT29, HCT-116, RKO, DLD1 and LoVo) by western blot and RHOT2 was confirmed to be expressed in all colon cancer cell lines (Figure 4E). To investigate the role of RHOT2 in the migration of colon cancer, RHOT2 interference fragments (si-RHOT2#1 and #2) were used in this study. The data in Figure 4F showed that the protein expression of RHOT2 was significantly decreased by si-RNAs, especially si-RHOT2#2. Then, we investigated the effect of RHOT2 in the migration of colon cancer cells. The results of the transwell assay showed that the migration ability of colon cancer cells was significantly increased by si-RHOT2 (Figure 4G, P < 0.05). Low expression level of RHOT2 markedly enhanced the migration ability of colon cancer cells (P < 0.05).

## Discussion

Here, we present the first comprehensive proteomics study to focus on LNM in patients with submucosal T1 CRCs. Through the MS-based proteomic technique, we developed a high-performance protein signature-based LNM prediction model to aid in decision-making regarding whether additional surgical resection is needed after endoscopic resection of T1 CRCs. The classifier was then validated in two different validation cohorts: a cohort with specimens removed by ESD and a prospective validation cohort. We also built an effective classifier using 5 proteins by their IHC score. We analyzed 221 T1 CRC patients and identified different molecular characteristics between patients with and without LNM. We also uncovered protein characteristics of mucinous colorectal adenocarcinoma.

Based on the T1 CRC proteomics dataset, we established a machine learning model to predict LNM. The protein signatures successfully stratified patients according to their risk of LNM. For construction of the protein signatures, in the training cohort, 105 candidate protein features were reduced to 55 potential predictors by examining the predictor-outcome association by shrinking the regression coefficients with the LASSO method. Based on 55 biomarkers, 10-fold cross-validation was applied to the training cohort, and an ROC curve was generated. The classifier achieved an AUC of 1.00 in the training cohort and was confirmed in two independent validation cohorts. The 55 protein markers achieved an AUC of 0.96 (95% CI, 0.917 to 1.000) in the ESD validation cohort and 0.933 (95% CI, 0.858 to 1.000) in the prospectively collected validation cohort. Next, 9 proteins with significant expression changes were selected to construct a new classifier. This classifier also had excellent performance for the identification of LNM when we assessed its calibration, discrimination, and clinical usefulness. We further performed IHC staining to confirm the expression patterns of the proteins and successfully predicted LNM with a single protein and built a predict model with 5 proteins.

Compared with previous studies, our protein-based predictive model has the following significant advantages besides high accuracy, since the ultimate goal of the project was to reduce additional surgery after endoscopic resection, we set up a cohort only containing EDS samples to validate our predictive model, and a prospective cohort was also used to validate the accuracy of the model. We further utilized immunohistochemistry to optimize our model for clinical use.

## Abbreviations used in this paper

LNM: lymph node metastasis
CRC: colorectal cancer
FFPE: formalin-fixed paraffin-embedded
VC1: validation cohort1
VC2: validation cohort2
IHC: immunohistochemistry
EVs: extracellular vesicles
miRNA: microRNA
mRNA: messenger RNA
TCGA: The Cancer Genome Atlas
AUC: area under the ROC curve
MS: mass spectrometry
KEGG: Kyoto Encyclopedia Genes and Genomes
GO: Gene Ontology
LASSO: least absolute shrinkage and selection operator

## Ethics approval

The present study was carried out comply with the ethical standards of Helsinki Declaration II and approved by the Institution Review Board of Fudan University Zhongshan Hospital (B2019-166). All patients signed the informed consent form.

## Data Availability Statement

All data generated or analysed during this study are included in the manuscript and supporting file; Source Data files have been provided for all figures. The raw data that support the findings of this study are available from the corresponding, CD, in iprox at https://111.198.139.98/page/PSV023.html;?url=1663254710170hUMW, reference number: IPX0003019000 upon reasonable request.

## Acknowledgements

This research was supported by Key Technologies Research and Development Program (2017YFA0505102, 2016YFC0901903, 2018YFA0507501, 2017YFC0908404); National Natural Science Foundation of China (31770886, 31972933, 91629301, 31700682); Shanghai Science and Technology Committee Project (19511121301); Clinical Research Plan of SHDC (SHDC2020CR5006).

## Author Contributions

Aojia Zhuang, PhD (Conceptualization: Lead; Data curation: Lead; Formal Analysis: Lead; Investigation: Equal; Methodology: Lead; Software: Lead; Visualization: Lead; Writing-original draft: Lead; Writing-review & editing: Lead)

Aobo Zhuang, MD (Conceptualization: Lead; Data curation: Equal; Formal Analysis: Equal; Investigation: Lead; Methodology: Equal; Resources: Supporting; Software: Supporting; Writing-original draft: Equal);

Zhaoyu Qin, PhD (Formal Analysis: Equal; Funding acquisition: Supporting; Methodology: Equal; Writing-review & editing: Supporting)

Dexiang Zhu, MD (Conceptualization: Supporting; Investigation: Supporting; Methodology: Equal; Writing-review & editing: Supporting);

Li Ren, MD (Investigation: Equal; Methodology: Equal);

Ye Wei, MD (Formal analysis: Equal; Resources: Supporting; Writing – review & editing: Equal);

Pengyang Zhou, MD (Investigation: Supporting; Methodology: Supporting;);

Xuetong Yue, MD (Investigation: Supporting; Methodology: Supporting;);

Fuchu He, Prof. (Conceptualization: Supporting; Methodology: Supporting; Project administration: Supporting; Resources: Supporting; Supervision: Equal);

Jianming Xu, Prof. Dr. (Conceptualization: Lead; Funding acquisition: Lead; Project administration: Lead; Resources: Lead; Writing-review & editing: Supporting);

Chen Ding, Prof. (Conceptualization: Lead; Funding acquisition: Lead; Investigation: Lead; Methodology: Lead; Project administration: Lead; Supervision: Lead; Writing-original draft: Equal; Writing-review & editing: Lead)

## Data Transparency Statement

This manuscript is an honest, accurate, and transparent account of our study. No important aspects of the study have been omitted and that discrepancies from the study as originally planned have been explained.

## Supplementary Figure Legends

**Figure 1- figure supplement 1.**
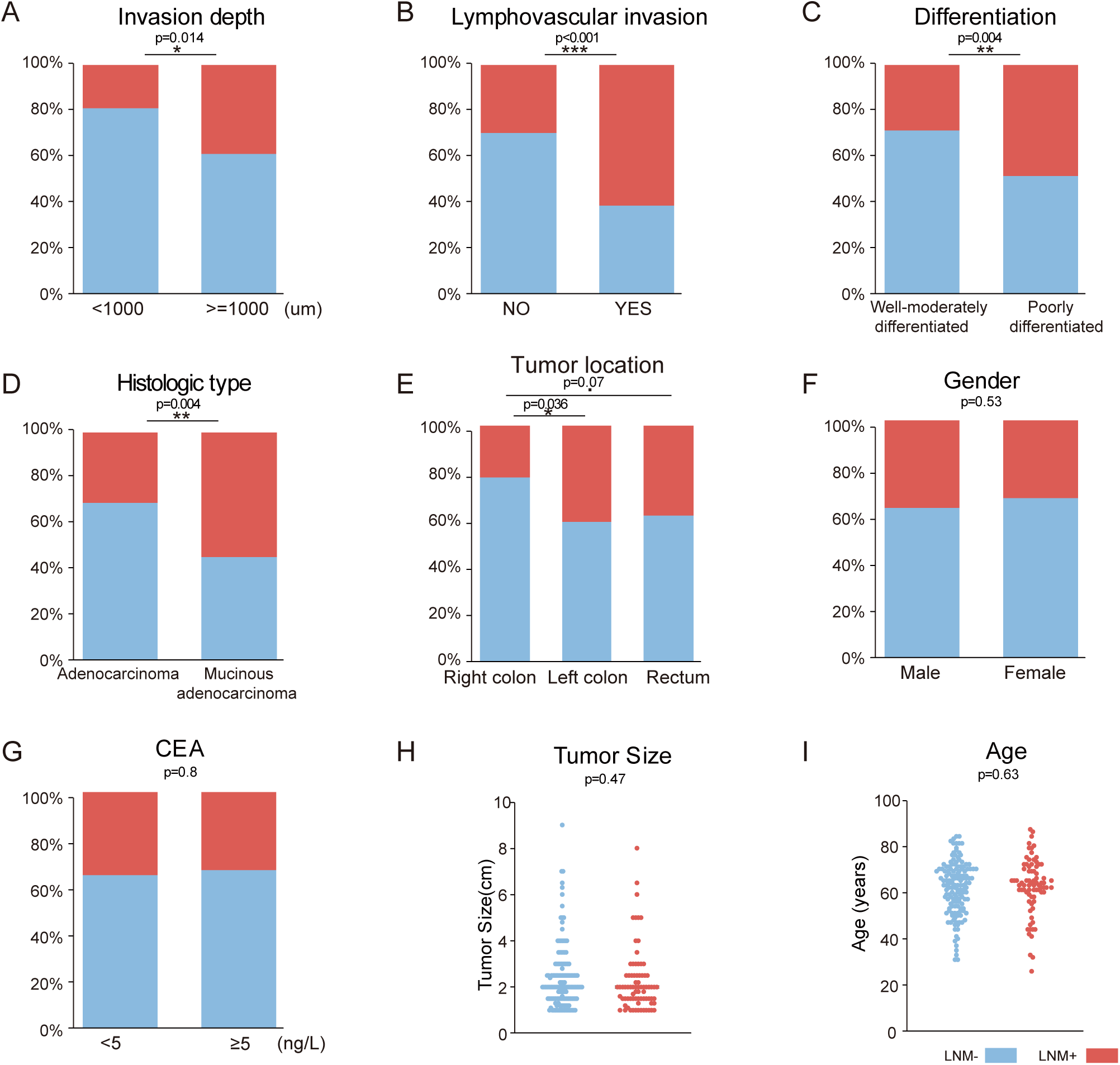
Association between clinical parameters and LNM. Related to Figure 1. A-I. The following clinical parameters were examined: invasion depth (A), lymphovascular invasion (B), differentiation (C), histologic type (D), tumor location (E), sex (F), CEA (G), tumor size (H), and age (I) (*p < 0.05; **p < 0.01, ***p< 0.001, chi-square test).

**Figure 1- figure supplement 2.**
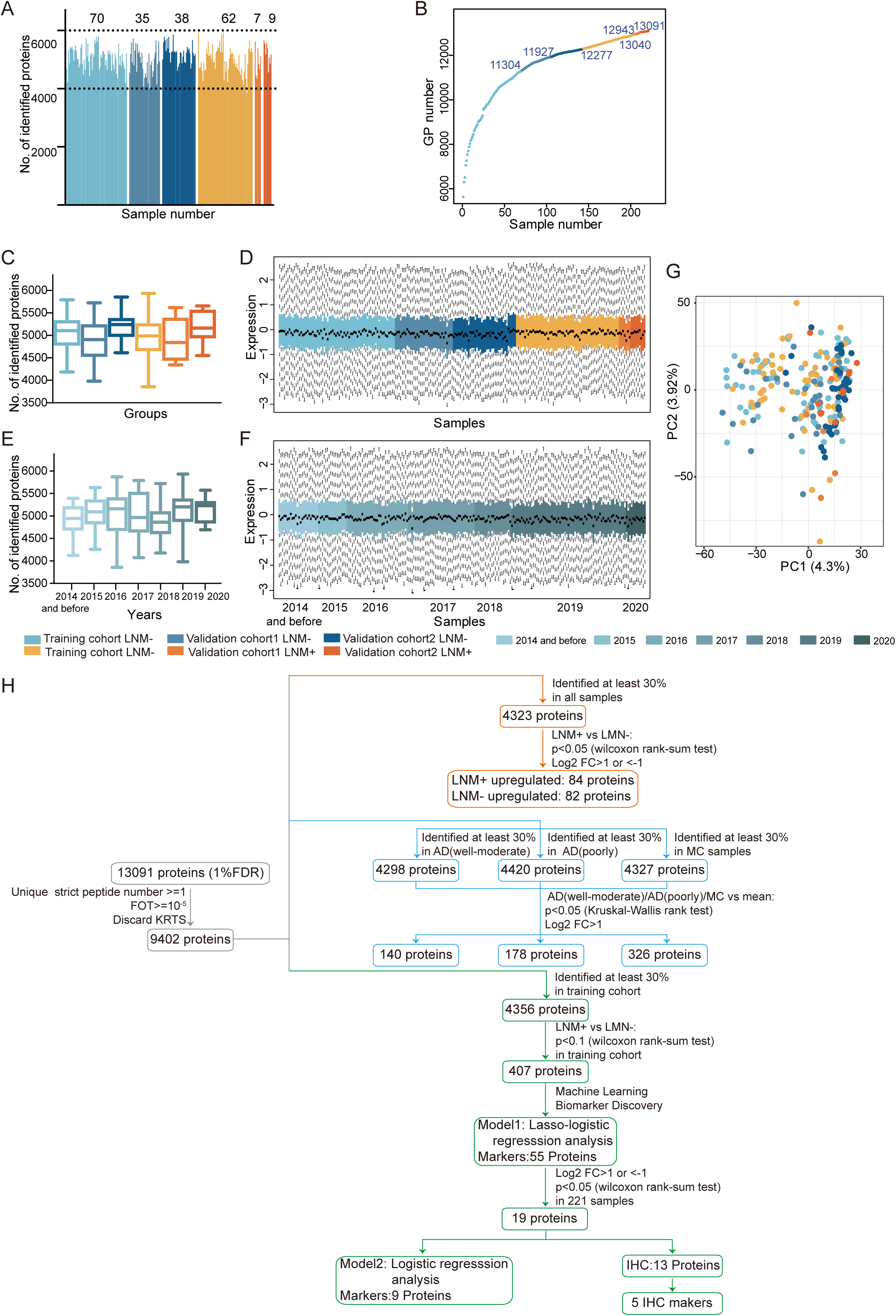
Quality assessment of proteomic data and summary of the analysis. Related to Figure 1. A. Total number of proteins quantified in 221 samples (LNM-positive samples from the training cohort (light orange), validation cohort 1 (orange), and validation cohort 2 (dark orange); LNM-negative samples from the training cohort (light blue), validation cohort 1 (blue), and validation cohort 2 (dark blue)). E. Cumulative number of proteins identified. B. Cumulative number of proteins identified. C&E. Box plots of the proteins identified in LNM-negative and LNM-positive samples from different cohorts (C)/sample collection time (E). LNM-negative samples from the training cohort (light blue), validation cohort 1 (blue), validation cohort 2 (dark blue); LNM-positive samples from the training cohort (light orange), validation cohort 1 (orange), and validation cohort 2 (dark orange). D&F. Distribution of log10-transformed iBAQ abundance of identified proteins in 221 proteome samples from different cohorts (D)/sample collection time (F) that passed quality control. G. Principal component analysis across different cohorts. The study samples were intermixed, suggesting a limited batch effect, and the quality control samples (QCs) clustered together, indicating good technical reproducibility. H. Proteomic datasets filtered at different levels for various statistical analyses.

**Figure 2- figure supplement 1.**
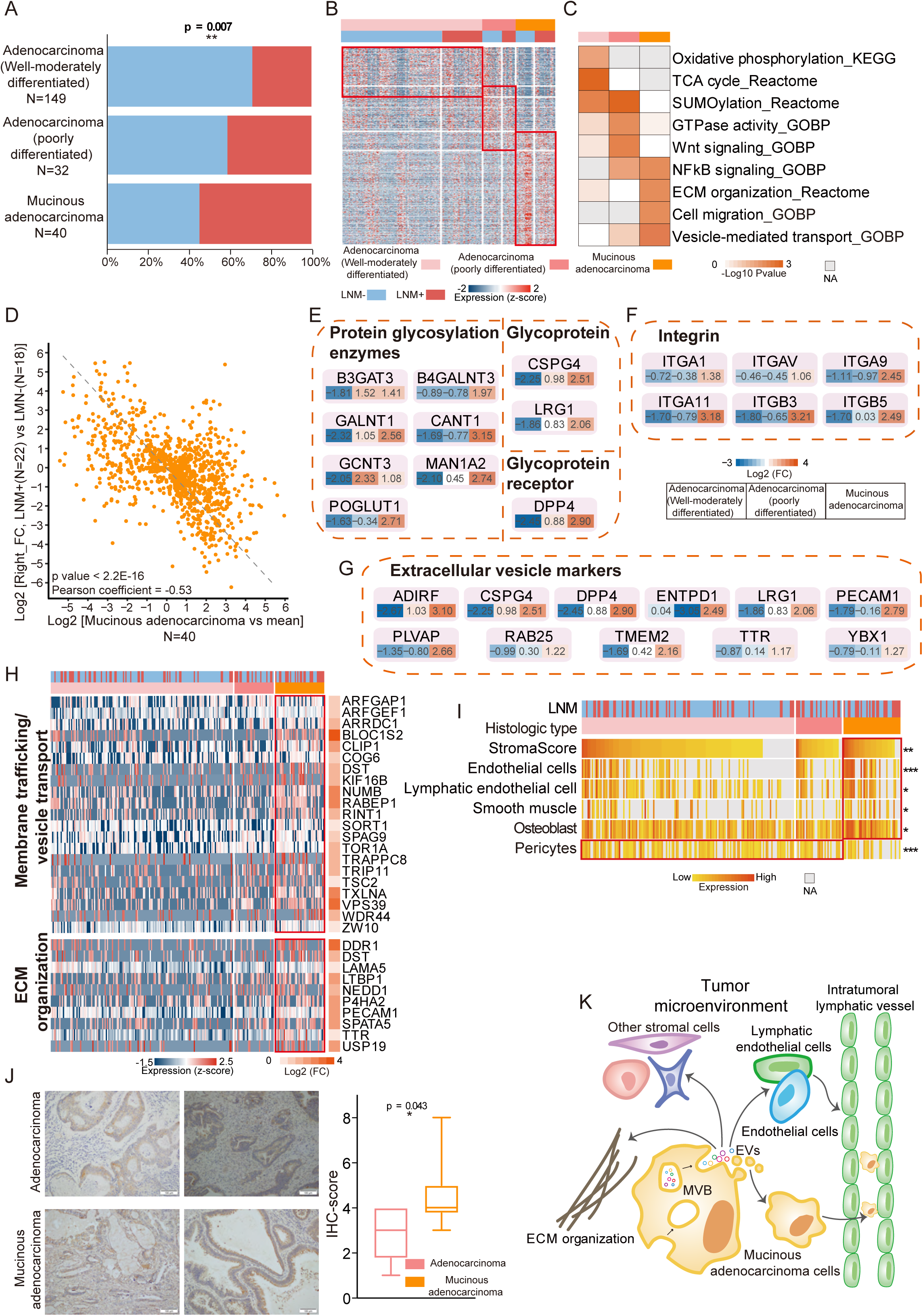
Protein differences by differentiation and histologic type and proteogenomic characteristics of mucinous colorectal adenocarcinoma. A. Bar graphs illustrating the relative proportion of LNM in well to moderately differentiated adenocarcinoma, poorly differentiated adenocarcinoma and mucinous adenocarcinoma. P values were calculated from the chi-square test. B. Upregulated proteins (proteins found in >30% of samples, a fold change more than 2 and p < 0.05 (Kruskal-Wallis test)) in T1 CRC with varying degrees of differentiation and different histologic types. Also see Figure 1- figure supplement 2H. C. Pathway enrichment (Reactome, KEGG, and GO) analysis showing upregulated pathways in T1 CRC with varying degrees of differentiation and different histologic types. D. Correlation of protein expression fold changes in LNM-positive/LNM-negative patients with histologic type (mucinous adenocarcinoma vs. the mean). E, F, G. Glycoprotein- and glycosylation-related protein (E), Integrins (F) and extracellular vesicle markers (G) were overexpressed in mucinous colorectal adenocarcinoma. H. Proteins in two pathways (membrane trafficking and/or vesicle transport and ECM organization) that were overexpressed in mucinous colorectal adenocarcinoma. I. Stromal score and signatures from xCell (*p < 0.05; **p < 0.01, ***p< 0.001, Kruskal-Wallis test). J. IHC staining of D2-40. Scale bar, 100 mm. Boxplots show the quantification of the IHC results. K. Model of LNM progression in mucinous colorectal adenocarcinoma.

**Figure 3- figure supplement 1.**
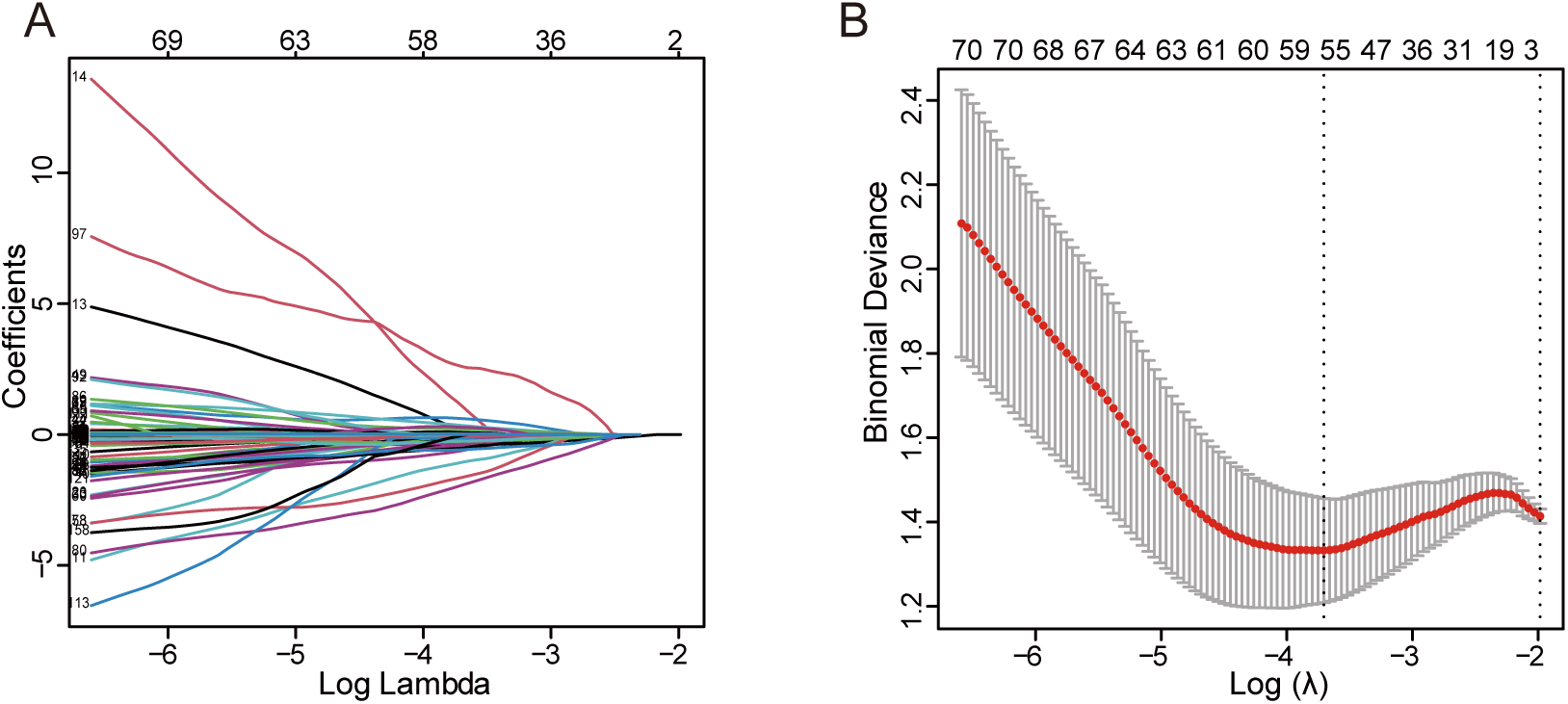
Details of the LASSO regression model and IHC staining of targeted proteins. Related to Figure 3. A. Tuning parameter (l) selection in the LASSO model via minimum criteria. B. LASSO coefficient profiles of the 105 texture features. A coefficient profile plot was produced against the log (l) sequence. A vertical line was drawn at the value selected using LASSO, where optimal l resulted in 55 nonzero coefficients.

**Figure 4- figure supplement 1.**
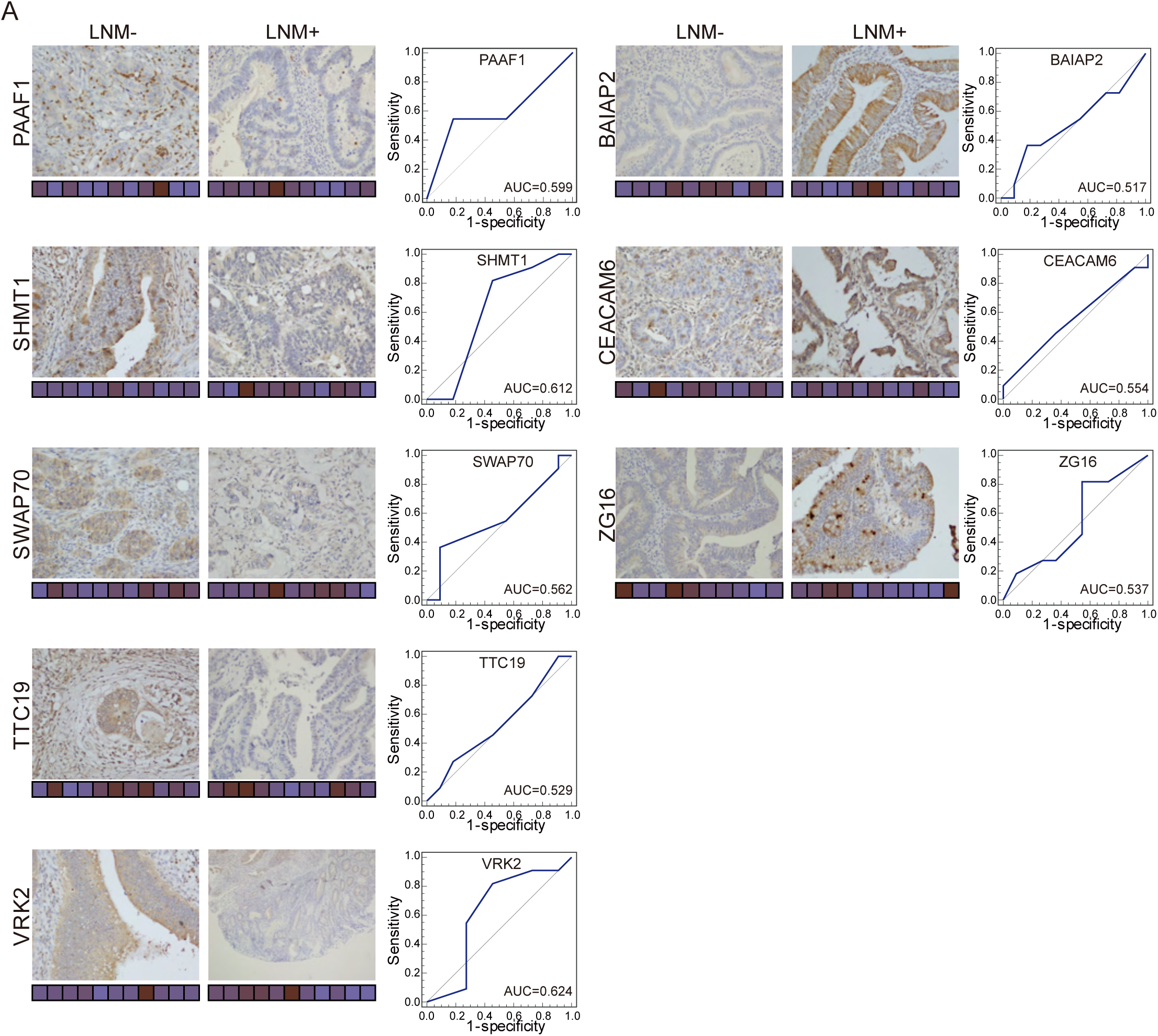
Immunohistochemical staining of targeted proteins. Related to Figure 4. T1 CRC samples from 22 patients were used to verify the abundance of PAAF1, SHMT1, SWAP70, TTC19 and VRK2 (left), and 21 samples were used to verify the abundance of BAIAP2, CEACAM6 and ZG16 (right). The scores that represent the product of the intensities and percentage of protein staining in the LNM-negative or LNM-positive samples are shown as a heat map.

Figure 1-source data1: Clinicopathologic Features

Figure1-source data2: All identified proteins

Figure 1-source data3: Filtered Proteomics Data

Figure 2-source data1: Log2 transformed proteomics data

Figure 2-source data2: Druggability based on the Drug Gene Interaction Database

Figure 2-source data3: Immune composition of T1 CRC from xCell

Figure 2-source data4: D2-40 Immunohistochemistry (IHC) staining score

Figure 3-source data1: Coefficients of 55 protein-markers and the LNM scores of samples using LASSO-logistic regression

Figure 3-source data2: predicting risk score for LNM of each patient

Figure 3-source data3: Coefficients of 9 protein-markers

Figure 4-source data1: Immunohistochemistry (IHC) staining score

Figure 4-WB source images

